# CEP76 is a critical regulator of male germ cell transition zone function and tail composition

**DOI:** 10.1101/2023.03.08.531794

**Authors:** Brendan J. Houston, D. Jo Merriner, G. Gemma Stathatos, Anne E. O’Connor, Alexandra M. Lopes, Donald F. Conrad, Mark Baker, Jessica E.M. Dunleavy, Moira K. O’Bryan

## Abstract

The transition zone is a specialised gate at the base of cilia/flagella, which segregates the ciliary compartment from the cytoplasm and strictly regulates protein entry. In this study, we have identified CEP76 as an essential germ cell transition zone protein, involved in the selective entry and incorporation of key proteins required for sperm function and fertility into the ciliary compartment and ultimately the sperm tail. In its absence sperm tails are shorter and immotile as a consequence of deficits in essential sperm motility proteins including DNAH2 and AKAP4, which accumulate at the sperm neck in the mutant. We demonstrate CEP76 is required for sperm tail fibrous sheath formation, outer dense fibre loading and axoneme stability in the principal piece and ultimately sperm motility. Finally, we identify that CEP76 dictates annulus positioning and composition, adding further evidence that the spermatid transition zone and annulus are part of the same functional structure.

## Introduction

Cilia and their organelle cousins, flagella, play essential roles in many cell types. Notably, a single modified motile cilium (a flagellum) projects from male gametes and is essential for fertility in sexually reproducing animals [1]. In eukaryotic cilia, the transition zone (TZ), also known as the ciliary gate, has emerged as an essential mediator of cilia development through its role in controlling protein entry into the ciliary compartment within which cilia/flagella develop [2–4]. The TZ develops immediately distal to the mature centriole, from which the core of the cilia, the axoneme, develops. The TZ allows for the selective transport of proteins into the cilium/flagellum and thus the establishment and maintenance of a unique ciliary microenvironment [5, 6]. The TZ is largely composed of sheets of Y-shaped structures that attach at one side to the developing axoneme, at a single point, and at the other side dock to the plasma membrane, at two points [7, 8]. While TZ composition, including Y-shaped linkers, is poorly understood, several genes that, when mutated, result in ciliopathies have been identified as core members of the TZ [9–12]. To enact protein and vesicle transport across the TZ and along the developing axoneme, cilia/flagella utilise a bi-directional transport process of intraflagellar transport (IFT) with the aid of the BBSome [13, 14]. In addition, the distal appendages of the centriole (known as transition fibres) play an essential role in protein trafficking into the ciliary compartment by acting as docking sites for cargoes prior to their entry through the TZ [15].

The sperm tail is a modified motile cilium and while it is assumed that the formation of the TZ and core of the axoneme will be similar to that which occurs in somatic cells, this is largely untested. Current models of the TZ are largely informed by data from primary cilia, or the somatic flagella of lower order species such as *Chlamydomonas* [16]. There is, however, emerging evidence to suggest differences in TZ structure, composition and transport machinery exist to meet the demands of different cilia/flagella sub-types (reviewed in [17]). As mammalian sperm contain accessory structures not seen in somatic cells, it is reasonable to predict that the male germ cell TZ is modified to selectively recruit fibrous sheath and outer dense fiber proteins into the ciliary lobe.

As with all motile cilia/flagella, sperm tail motility is dictated by the function of the axoneme – a 9+2 microtubule-based structure that runs the length of the tail [18, 19]. The sperm tail is composed of two major sections: the midpiece, which houses the mitochondria; and the principal piece, wherein all glycolytic enzymes are anchored to a structure called the fibrous sheath [20–23]. Both the midpiece and principal piece compartments also contain outer dense fibres – which are circumferential to the axoneme microtubules and act to protect the sperm tail from shearing forces [24, 25]. In the principal piece, the longitudinal columns of the fibrous sheath replace the outer dense fibres 3 and 8 and are linked by circumferential ribs [26, 27]. As per the axoneme, the outer dense fibres and fibrous sheath are formed within the ciliary compartment and as such their component proteins must transition through the TZ [24]. While this may suggest the requirement of additional factors to aid in the selective transport of sperm-specific proteins via the TZ into the ciliary compartment, no such proteins of this function have been identified.

In addition, towards the end of sperm tail development, the annulus, a septin-based ring that is thought to be attached to, or physically a part of the TZ, migrates distally along the sperm tail to define the junction of what will ultimately become the midpiece and principal pieces of the sperm tail [28, 29]. This annulus is thought to be a diffusion barrier between the sub-compartments of the tail, and its loss is often associated with a sharp bending at the midpiece-principal piece junction, poor sperm motility and ultimately male infertility [28, 30]. At a similar time to annulus migration, but by a seemingly independent process [28, 31, 32], the membrane attached to the basal body is pulled distally to the annulus, thus exposing a portion of the axoneme (surrounded by the outer dense fibres) to the cytoplasm. This process allows cytoplasmic mitochondria to be loaded onto the sperm tail to form the mitochondrial sheath of the midpiece [24, 33].

Despite our descriptive understanding of a number of these processes, the mechanisms of sperm tail development are still largely unknown. It is currently unclear how sperm-specific proteins, including components of the accessory structures, are selectively transported via the TZ into the ciliary compartment. In working to address this, we identified the previously unexplored centriole gene *CEP76* as being essential for male fertility in men as highlighted by a missense mutation in an infertile man with azoospermia (Figure S1; [34, 35]). CEP76 has been suggested to play a role in centriole duplication in somatic cell lines through interaction with the centriole protein CP110 [36, 37], but nothing was known about its role in male fertility. *CEP76* expression is testis enriched [38] and CEP76 has been localised to centrioles in human spermatozoa and somatic cells [37, 39]. CEP76 has two putative functional domains (Figure S1): a C2 domain that is predicted to function in ciliary membrane targeting, and a transglutaminase domain that is predicted to interact with tubulins in the axoneme or the TZ [40]. It is hypothesised based on a large TZ protein comparative study, that these domains cooperate to play a role in Y-shaped linker function (Zhang and Aravind, 2012) and likely TZ function. Thus, we aimed to define the role CEP76 plays in male fertility.

Within this study we identify CEP76 as an essential male fertility gene, with a unique role in TZ function and the selective entry of key motility proteins into the developing flagellum. Knockout males were sterile and produced sperm with a variety of structural defects and the abnormal accumulation of key proteins at the sperm neck, consistent with an inability to pass through the TZ during the process of tail development. Consequently, formation of the mitochondrial sheath was significantly impaired in sperm from *Cep76* knockouts. Collectively, these data identity CEP76 as the first known germ cell-specific regulator of transition zone function, which is required for male fertility in mammals.

## Results

### *Cep76* is a spermatid enriched mRNA

Consistent with its putative role in male fertility, mouse *Cep76* is most highly expressed in the testis (Figure S2A) with lower levels also detected in the brain. To define which germ cell types *Cep76* is expressed in, we investigated its expression across the establishment of the first wave of mouse spermatogenesis (Figure S2B). *Cep76* mouse testis expression levels were relatively consistent until day 18 of age, and then notably increased at day 20 of age, coincident with the beginning of spermiogenesis (first appearance of round spermatids). In agreement, single cell RNA sequencing of mouse germ cells revealed that *Cep76* expression is considerably elevated in step 6-9 spermatids (Figure S2C), at the time when the core of the sperm tail is formed. We tested multiple antibodies to investigate CEP76 localisation in male germ cells but found all were non-specific (data not shown).

### *Cep76* is required for male fertility in mice

To directly test the hypothesis that CEP76 is required for sperm tail development and thus male fertility, we generated a knockout mouse model using CRSIPR/Cas9 technology. We removed exon 3 of the mouse gene (Figure S1E), which resulted in a premature stop codon in exon 4 of *Cep76*, which produces one protein coding transcript (ENSMUST00000097542.3). This transcript encodes a protein of identical length (659 AAs, predicted molecular mass of 74.3 kDa) and 97.7% identity (Figure S1B) to the human *CEP76* principal isoform. Additionally, sequence comparison revealed that the amino acid affected in the infertile man is a highly conserved residue from men to zebrafish (Figure S1B). *Cep76* knockout was confirmed via qPCR (Figure S1F).

While knockout males were free of overt systemic disease and displayed normal mating behaviour, they were sterile (Figure 1A). Knockout females were fertile (not shown). Body, testis, and epididymis weights (Figures 1B, C and D) of *Cep76* knockout males were equivalent to wild type males. While the daily sperm production of *Cep76* knockout males was similar to wild type counterparts (Figure 1E), the number of sperm within their epididymides was reduced to approximately 35% of wild type levels (Figure 1F; *p* < 0.0001). The reduced sperm number in the *Cep76* knockout epididymis, in the absence of a reduction in testis weight and daily sperm production was due to partial spermiation failure, as evidenced histologically in stage IX tubules, wherein sperm were retained within the seminiferous epithelium (Figure 1J). With the exception of spermiation failure, spermatogenesis appeared normal at a light microscopic level in *Cep76* knockouts (Figure 1H). Similarly, epididymal histology was comparable between genotypes (Figures 1K and L).

**Figure 1.**
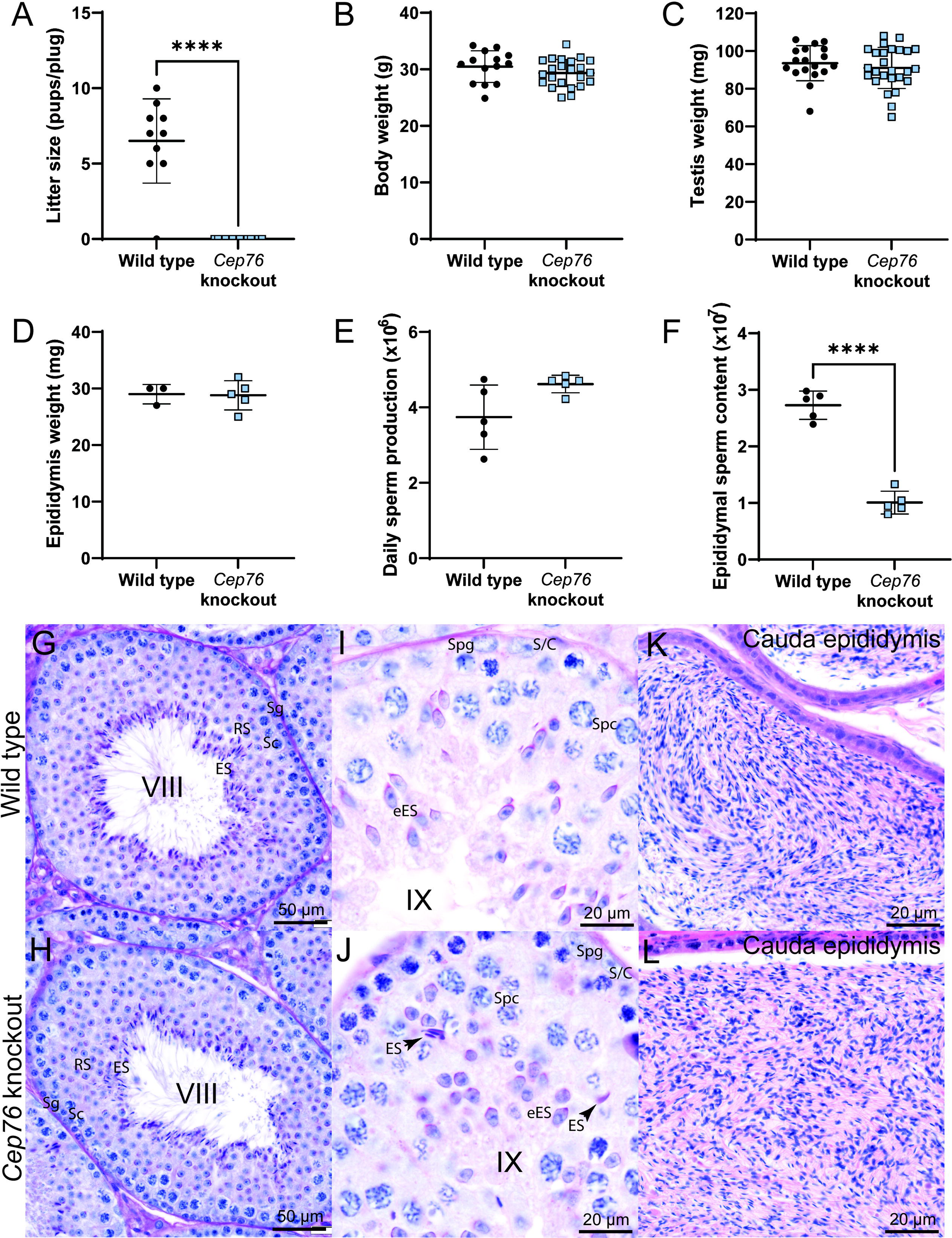
CEP76 is essential for male fertility. Wild type versus *Cep76* knockout data for A. Litter size. B. Body weight. C. Testis weight. D. Epididymis weight. E. Daily sperm production per testis. F. Epididymal sperm content. **** *p* < 0.0001. G, I. Histology of wild type and H, J. *Cep76* knockout testis sections of stage VIII and stage IX tubules, respectively. Arrows in J point to retained spermatids as evidence of spermiation failure. Sg = spermatogonia, Sc = spermatocyte, RS = round spermatid, ES = elongating spermatid, S/C = Sertoli cell. K, L. Cauda epididymis sections are shown for wild type and *Cep76* knockout. Scale bars are noted to equal 50 or 20 µm.

#### CEP76 is required for normal sperm morphology and motility

Analysis of sperm morphology via light microscopy revealed overt defects in sperm from knockout males (Figure 2A). Total sperm tail and midpiece length (measured by mitochondrial sheath length) of sperm from *Cep76* knockout males was significantly shorter (by 15% and 25%, respectively) than those from wild type males (Figures 2B and C; *p* < 0.0001), suggesting a fundamental role of CEP76 in tail assembly within the sperm ciliary compartment that is consistent with a role in TZ function and mitochondrial loading. In agreement with the defects in sperm morphology, computer assisted sperm analysis revealed a striking reduction in the percentage of sperm from knockouts displaying basic twitching motility (Figure 2A; 6% in knockout versus 80% in wild type, *p* < 0.0001) and the virtual absence of forward, progressively motile sperm in comparison to wild type males (Figure 2B; 0.4% in knockout versus 42.4% in wild type, *p* < 0.0001). The small population of sperm from *Cep76* knockout males that were classified as motile were seen to be simply twitching on the spot.

**Figure 2.**
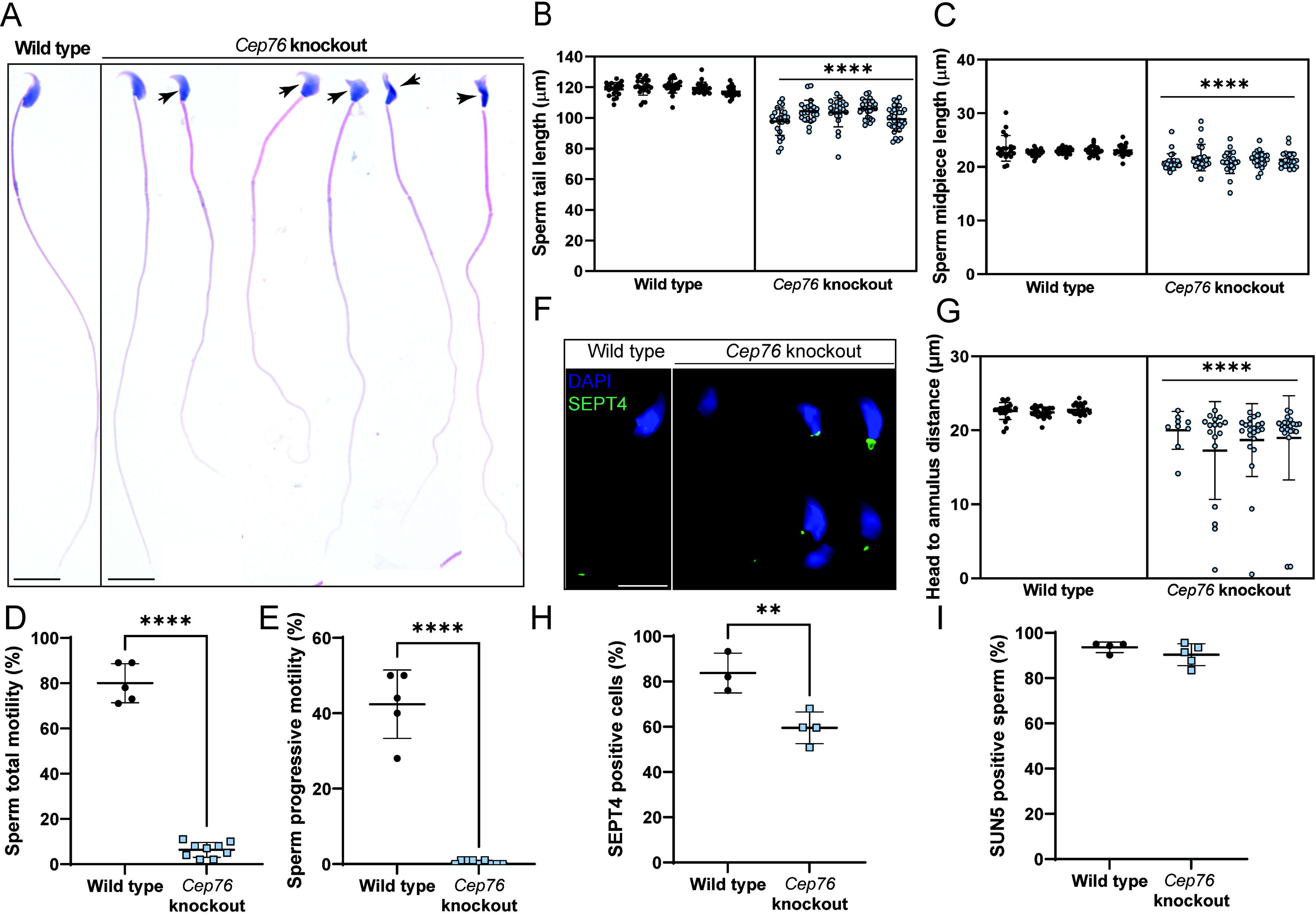
CEP76 is required for normal sperm morphology, motility and tail length. Wild type versus *Cep76* knockout data for A. Total and B. Progressive sperm motility as assessed by computer assisted sperm analysis. C. Sperm morphology – arrows denote abnormally shaped sperm heads. D. Total sperm tail length. E. Sperm midpiece length as measured via MitoTracker staining. F. Sperm annulus staining (SEPT4). Scale bar = 5 µm. G. Sperm annulus migration distance. H. Incidence of SEPT4 staining in sperm. I. Incidence of sperm SUN5 staining. ** *p* < 0.01, **** *p* < 0.0001.

To examine the cause of the reduced mitochondrial sheath length in the absence CEP76, we measured the distance between the base of the sperm head and the annulus as marked by SEPT4 staining (Figure 2F, G). Measurements revealed a significantly shorter distance between the annulus and nucleus in sperm from *Cep76* knockout males relative to wild type (18.6 µm in knockout versus 22.6 µm in wild type; *p* < 0.0001). In addition, the portion of sperm with SEPT4 positive staining (Figure 2H) was significantly reduced in the absence of CEP76 (60% in knockout versus 84% in wild type, *p* = 0.0095), suggesting poorly formed annuli.

Light microscopy and a visual examination also revealed defects in sperm head shape (Figure S3A) in at least 40% of sperm from knockouts compared to 5% in wild type (*p* < 0.0001). We also observed a significant increase in sperm with abnormal acrosomes from *Cep76* knockout males (35% in knockout versus 3% in wild type, Figure S3D; *p* < 0.0001). In addition, we analysed sperm head morphology using a nuclear shape analysis software. This revealed a significantly lower proportion of the knockout population classified as normal (Cluster 1), while there were significant increases in sperm with slightly

(Cluster 2) and largely (Cluster 4) abnormal head shapes (compared to wild type). This analysis revealed that ∼70% of sperm heads from *Cep76* knockout males had abnormal nuclear morphology (Figure S3E; *p* < 0.0001). In addition, there was a 2-fold increase in sperm decapitation in samples from knockout males, indicating that the CEP76 plays a role in establishing patency of the head-tail coupling apparatus (HTCA) (Figure S3C, *p* < 0.0001).

Collectively these results reveal CEP76 is required for the production and release of normal numbers of functional sperm. Shorter sperm tails and annulus defects suggest CEP76 plays a role in TZ function. Defects in nuclear morphology and a weakened HTCA are supportive of a role for CEP76 in manchette function and/or acrosome formation and in the fortification of the neck region (a derivative of the basal body) late in spermiogenesis. At a function level, sperm from *Cep76* knockout males are unable to reach the site of fertilisation due to highly impaired sperm motility.

### CEP76 is required for normal sperm flagella and HTCA ultrastructure

In order to define the origins of the motility and structural defects, we investigated sperm ultrastructure using transmission (TEM) and scanning electron microscopy (SEM) on isolated epididymal sperm (Figures 3, 4, Figure S4). At the midpiece level, the axoneme ultrastructure appeared superficially normal, bearing all outer dense fibres and the 9+2 microtubule formation with dynein arms in sperm from wild type and knockout males (Figure 3A, B, respectively). In *Cep76* knockout sperm, however, a build-up of mitochondria, and membranes was seen at midpiece level cross-sections (Figure 3C). Longitudinal sperm tail sections of the midpiece from knockout males additionally revealed abnormal mitochondria with enlarged spacing within mitochondrial matrixes (Figure 3H) and notable mitochondria aggregation (Figure 3I – arrow).

**Figure 3.**
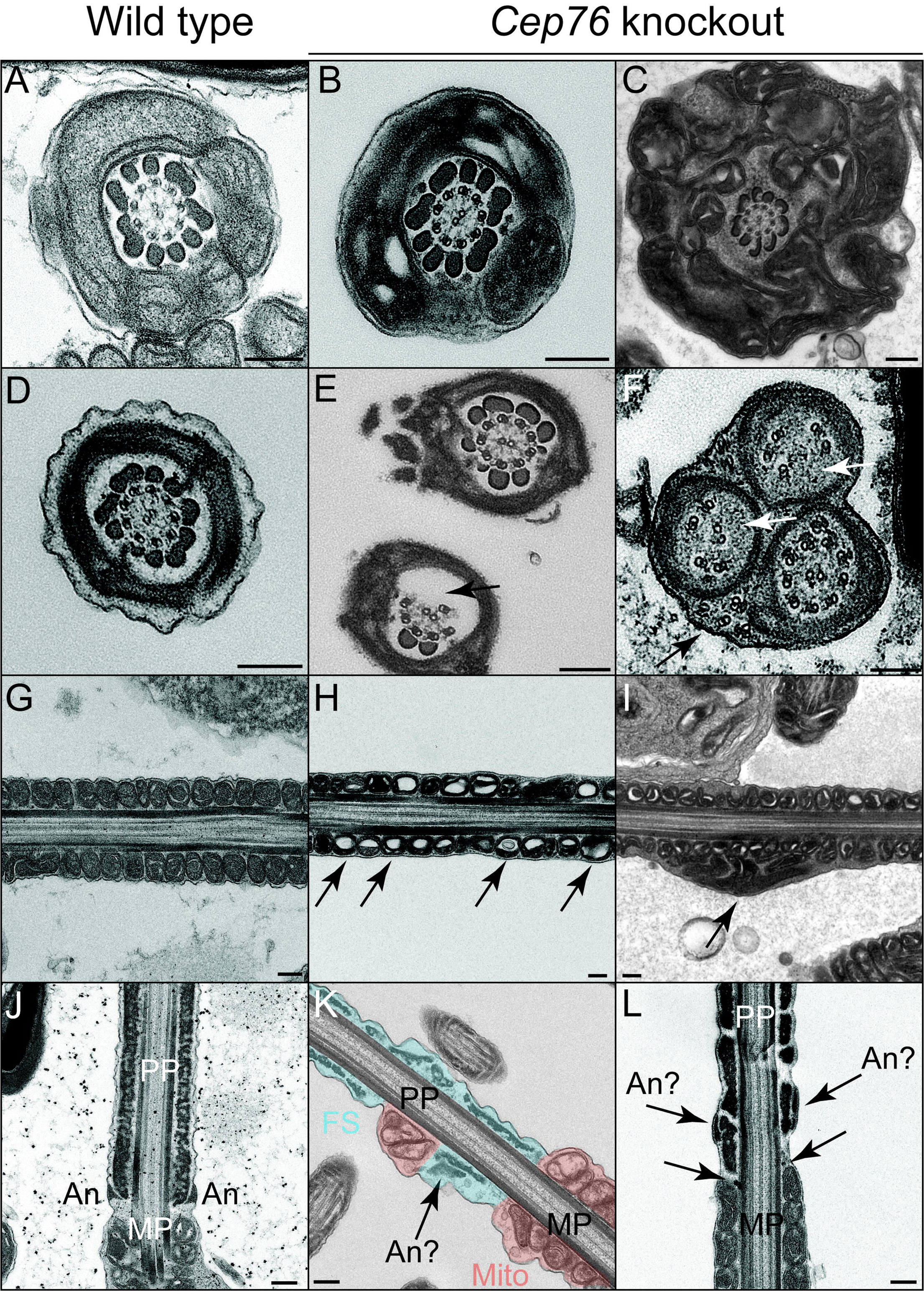
CEP76 is required for accessory structure assembly and normal axoneme ultrastructure. Wild type sperm are shown in panels A, D, G and J; sperm from *Cep76* knockouts are shown in panels B, C, E, F, H, I, K and L. **Top row** (A, B, C) – sperm midpiece cross-sections highlighted a broadly normal axoneme in both genotypes, with evidence of mitochondrial and membrane aggregation in knockout sperm. **Second row** (D, E, F) – sperm principal piece cross-sections highlighted the absence of some microtubule doublets and outer dense fibres (arrow, E), and the displacement of some microtubule doublets (black and white arrows, F) in knockout cells. **Third row** (G, H, I) – midpiece longitudinal sections highlighted abnormal mitochondrial morphology (arrows, H) and aggregation (arrow, I) in knockout cells. **Bottom row** (J, K, L) – longitudinal sections of the midpiece-principal piece boundary revealed abnormal annulus formation (arrows point to predicted annulus structures, K, L) and the consequential mixing of mitochondria and fibrous sheath structures in knockout cells (K). MP = midpiece, PP = principal piece, An = annulus, FS = fibrous sheath, Mito = mitochondria. Scale bars = 200 nm.

**Figure 4.**
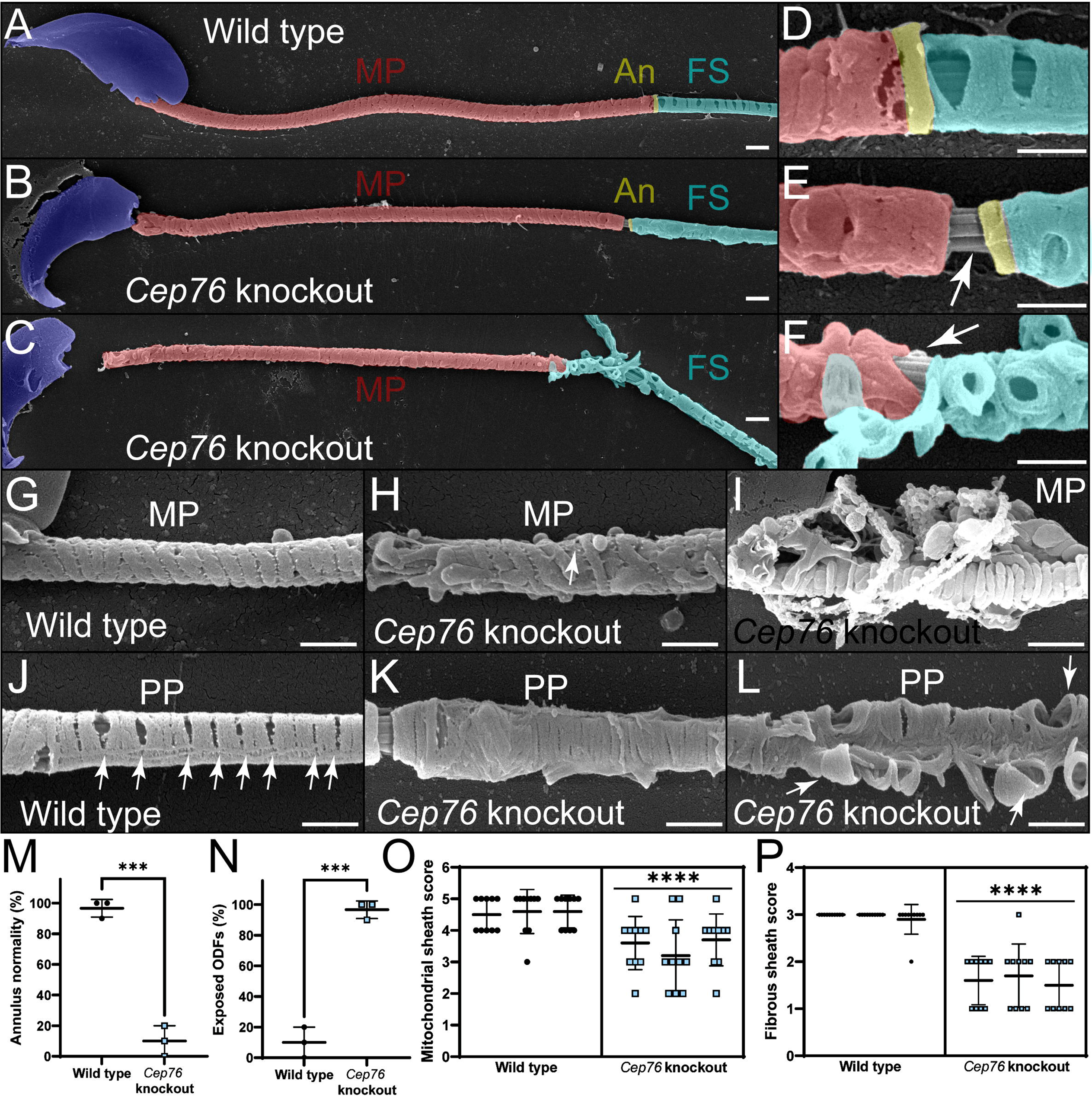
Annulus, mitochondrial sheath and fibrous sheath formation are impaired in the absence of CEP76. Scanning electron microscopy on membrane stripped sperm revealed defects in accessory structures in knockout sperm (B, C). MP = midpiece, An = annulus, FS = fibrous sheath. Sperm nuclei are overlayed in purple, mitochondria in red, annuli in yellow and fibrous sheaths in blue. High power images of the midpiece-principal piece boundary (D, E, F) revealed an abnormal annulus region in knockout sperm and the core axoneme being exposed (arrows). Wild type mitochondria were helically arranged with morphology normal (G), whereas mitochondria were poorly arranged (H) or aggregated (I) in knockout cells. Fibrous sheath deposition was highly abnormal in the absence of CEP76 (C, K, L) and appeared to be unanchored in knockout cells. Additionally, the slits present in wild type sperm (J, arrows) were missing from knockout cells. Scale bars = 1 µm. These defects were quantified (M-P). M. Annulus normality assessment. N. Prevalence of exposed outer dense fibres (ODFs). O. Mitochondrial sheath normality assessment (5 = best, 1 = worst). P. Fibrous sheath normality assessment (3 = best, 1 = worst). *** *p* < 0.001, **** *p* < 0.0001.

Investigation of the axoneme ultrastructure at the principal piece level revealed absent outer dense fibres and microtubule doublets in many sperm tails from knockout males (Figure 3D-F), suggesting this compromise, which was absent at the midpiece level, may have been influenced by fibrous sheath assembly. In addition, we observed the abnormal formation of the annulus, again suggesting an origin of poor TZ formation. Specifically, in sperm from *Cep76* knockout males the annulus was not clearly distinguishable from the fibrous sheath (Figure 3K), or not identifiable (Figure 3L). In addition, we observed rare examples of mitochondria incorporated within the principal piece in *Cep76* knockout male sperm (Figure 3K), but never in sperm from wild type controls.

Examination of the sperm neck region by TEM (Figure S4D-F) revealed a number of abnormal structures, including imperfect capitulum structures and a build-up of mitochondria and ectopic vesicles in sperm from *Cep76* knockout males. Further, an excess of what we predicted to be granulated bodies was seen throughout the cytoplasm (Figure S4E). Granulated bodies are transported into the ciliary compartment to form the outer dense fibres which is continuous with the HTCA [42]. This finding strongly supports a role for CEP76 in the selective entry of proteins and vesicles into the ciliary compartment during spermiogenesis and, by extension, in building of the ODF and the fortification of the HTCA. Of relevance, we did not observe a significant difference in the proportion of sperm positive for SUN5 staining, a protein required for the fortification between the capitulum and basal plate [43], between genotypes, i.e., the junction with the nuclear membrane (Figure 2I; *p* = 0.26). As this structure is proximal to where the TZ exists, this data is consistent with a role for CEP76 in the TZ and the assembly of more distal regions of the tail.

### CEP76 is required for the assembly of sperm tail accessory structures and annulus migration

#### Annulus

Scanning electron microscopy of sperm from *Cep76* knockout males further emphasised the poorly formed annulus and accessory structures in the midpiece and principal piece (Figure 4). In sperm from wild type males, the annulus was appropriately positioned at the junction between the mitochondrial sheath of the midpiece and the fibrous sheath of the principal piece (Figure 4A, D). In sperm from *Cep76* knockout males, while the annulus was often associated with the fibrous sheath, it was rarely found aligned at the distal border of the mitochondrial sheath (Figure 4B, C, E, F). As a result, a region of exposed ODFs was evident between the mitochondrial sheath and the annulus or fibrous sheath. Quantification of the defects in annulus positioning (Figure 4M) revealed a normal positioning and structure of the annulus in only 10% of sperm from knockouts males compared to 95% of sperm from wild type males (*p* = 0.0002). Consequently, we quantified the incidence of ODF-exposed regions at the distal midpiece (Figure 4N). This revealed that ODFs were exposed in 97% of sperm from *Cep76* knockouts but only 10% of sperm from wild type (*p* = 0.0002). In many cases, the annulus was found embedded within the proximal region of the fibrous sheath (e.g., Figure 4K) rather than at the junction of the midpiece, or it was not clearly observed due to large deformations in the fibrous sheath (Figure 4F). Due to the predicted role of CEP76 in TZ function, we anticipate this is due to a defect in TZ/annulus migration rather than the fibrous sheath assembling too far proximally.

### Mitochondria

As above, mitochondrial packing in epididymal sperm from *Cep76* knockouts was irregular and characterised by misaligned and poorly compacted mitochondria along the mitochondrial sheath (Figure 4H versus 5G). Mitochondrial aggregation defects identified via TEM were confirmed via SEM and were characterised by collections of poorly formed mitochondria and a build-up around the midpiece (Figure 4I). Scoring of mitochondrial sheath normality (as defined in the methods; Figure 4O) revealed a significant reduction in quality in sperm from *Cep76* knockout males compared to wild type (3.5 in knockout versus 4.6 in wild type; *p* < 0.0001).

### Fibrous sheath

Finally, SEM revealed that fibrous sheath formation was notably impaired in the absence of CEP76 (Figures 4C, F, K, L). In wild type sperm, the fibrous sheath is comprised of two longitudinal columns linked by circumferential ribs. Gaps in the fibrous sheath were evident at regular intervals (Figure 4D, J). In the absence of CEP76, however, gaps/slits between circumferential ribs were rare (Figure 4K) and in many cases the ribs appeared to be unanchored and projected from the body of the tail (Figure 4L). Notably, the proximal portion of the fibrous sheath appeared to be the most disorganised (e.g., Figure 4C). Fibrous sheath normality was scored (as detailed in the methods; Figure 4P), revealed a highly significant reduction in fibrous sheath quality in sperm from *Cep76* knockouts (1.6 in knockout versus 2.9 in wild type; *p* < 0.0001).

Collectively, these data reveal CEP76 is a key determinant in the development of multiple aspects of sperm tail development, including the fibrous sheath, annulus, and positioning and the mitochondrial sheath.

### Loss of CEP76 leads to aberrant sperm composition

To explore the hypothesis that CEP76 is involved in the selective entry of proteins into the ciliary lobe at a molecular level, we performed quantitative mass spectrometry on sperm from the cauda epididymis of *Cep76* knockout and wild type males (Table 1A). Doing so identified 32 differentially expressed proteins in sperm from knockout males (13 up-regulated and 19 down-regulated), including multiple mitochondrial and apoptotic proteins. This difference in mitochondrial protein content is consistent with the abnormal accumulation of mitochondria around the sperm tail as detailed above. In agreement with the observed reduced tail length, alpha tubulin content was significantly reduced in sperm from *Cep76* knockout males. In a separate analysis, and to account for differences in tail length, the content of known (or predicted) sperm tail proteins was normalised to alpha tubulin. As shown in Table 1B, following this normalisation, 14 proteins were identified as significantly altered in sperm from *Cep76* knockout males compared to wild type including several that are essential for sperm tail function and male fertility. They included: axonemal protein DNAH2 (1.65-fold wild type levels, *p* = 0.0406), the actin-based motor protein myosin 9 (3.5-fold wild type levels, *p* = 0.0174) and AKAP3 and AKAP4, which are major components of the fibrous sheath [22] (1.36-fold and 1.46-fold wild type levels, *p* = 0.032 and *p* = 0.0498, respectively).

To investigate this counterintuitive increase in several motility proteins, we defined the localisation of DNAH2 and AKAP4 proteins in sperm from *Cep76* knockout males using immunofluorescence (Figure 5). As expected, in wild type sperm, the dynein arm protein DNAH2 was uniformly localised throughout the midpiece and principal piece of the tail (Figure 5A) [44, 45]. In contrast, while DNAH2 was found within the sperm tail in many sperm from *Cep76* knockouts, it was notably accumulated in the neck region. In other sperm, DNAH2 was seen nearly exclusively localised to the neck and appeared to be almost absent from the tail (Figure 5A). Specifically, 22% of sperm from *Cep76* knockout males displayed an accumulation of DNAH2 in the neck region compared to 5.1% of sperm from wild types (4.3 x wild type levels, *p =* 0.0006; Figure 5C). Tail pixel intensity analysis (per area) revealed a 25% reduction in DNAH2 content across the entire flagellum (*p* < 0.05; Figure 5E).

**Figure 5.**
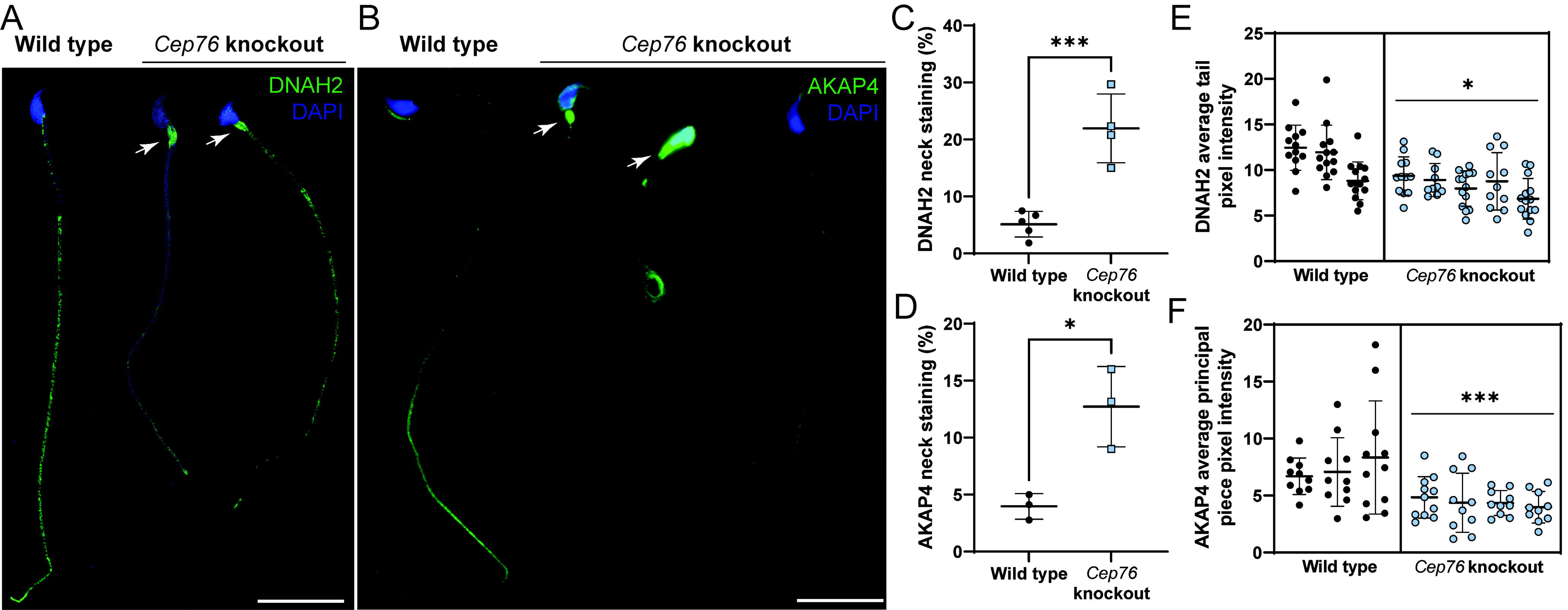
CEP76 is required for the loading of essential motility and fibrous sheath proteins into the sperm tail. Wild type versus *Cep76* knockout data for A. DNAH2 localisation in cauda epididymal sperm. B. AKAP4 localisation in cauda epididymal sperm. Scale bars = 20 µm. Arrows point to accumulation of DNAH2 or AKAP4 in the neck region of sperm. The number of sperm with this neck localisation was quantified, as shown in C and D for DNAH2 and AKAP4, respectively. The average tail pixel intensity (per area) of DNAH2 and AKAP4 were quantified and are shown in E and F. *** *p* < 0.001, * *p* < 0.05.

Also as expected for a component of the fibrous sheath, AKAP4 was primarily localised to the principal piece sperm from wild type (Figure 5B). As above, in *Cep76* knockout sperm, AKAP4 accumulated in the neck region in 13% of sperm compared to 4% in sperm from wild type mice (Figure 5D).

Quantification of neck AKAP4 staining revealed 3.1-fold levels in *Cep76* knockout sperm relative to wild type (Figure 5D). An analysis of pixel intensity revealed a significant, 40% reduction in AKAP4 content (per area) within the principal piece of sperm from *Cep76* knockout males compared to wild type (*p* < 0.0001; Figure 5F).

Collectively, these data underscore an essential role for CEP76 in TZ function and the regulated entry of key sperm tail proteins into the ciliary compartment and sperm tail development.

### CEP76 is not required for manchette formation and migration

SEM confirmed the magnitude of head deformation in sperm from *Cep76* knockout males (Figure S4A-C). The manchette is critically involved in shaping the proximal half of the sperm nucleus and further acts as a transport freeway for protein delivery to the basal body and into the sperm tail [24]. Despite the highly irregular sperm head shape, the formation of the manchette appeared normal in the absence of CEP76 as identified by alpha tubulin staining of isolated germ cells (not shown) and testis sections (Figure S3F). Electron microscopy reinforced that manchette structure was overtly normal (not shown).

### CEP76 is required for the maintenance of centriole number in male germ cells

CEP76 has previously been linked to a role in the suppression of centriole duplication in 293T cells [37], thus raising the possibility of a similar role in germ cells. To explore this hypothesis, we stained purified round and elongating spermatids with the centriole marker centrin (Figure S6G, H) and quantified the number of centrioles per cells (Figure S6I). As expected, two distinct centrin foci were observed in 84% of wild type spermatids and only 1% of wild type cells exhibited 3+ centrin foci. By contrast, in spermatids from *Cep76* knockout males, 18% of cells possessed 3 or more centrioles structures (*p* = 0.0002) and only 70% of cells contained 2 centrin foci (p = 0.0012). These data confirm a role for CEP76 in germ cell centriole duplication suppression. Despite this finding, sperm from *Cep76* knockouts contained only single tails, i.e., centriole overduplication did not lead to multiple basal bodies and axoneme growth.

## Discussion

Building a sperm tail is a complex and multistep process, requiring the coordinated action of multiple protein and organelle transport processes (reviewed in [24, 46]), and is absolutely essential for male fertility. The spermatid centriole is inherited during the process of meiosis. It subsequently duplicates and matures to give rise to the basal body that docks to both the plasma and nuclear membranes [47, 48]. From this structure, the sperm tail, a modified cilium, forms through the selective entry of proteins into the ciliary compartment via the TZ that sits at the junction between the cytoplasm and the ciliary compartment. Assuming that cilia formation during germ cell development is analogous to primary cilia wherein TZ function has been studied, all component proteins for the axoneme, outer dense fibers and fibrous sheath must be selectively transported through the TZ. In this study, we establish CEP76 as the first protein shown to play a male germ cell specific role in the TZ and demonstrate that CEP76 is required for the development of functionally competent sperm. We show CEP76 is essential for the efficient transport of AKAP4 proteins into the flagella compartment, underpinning normal fibrous sheath development. We also show that it facilitates the incorporation of tubulin into the tail, and that an absence of CEP76 leads to short sperm tails. Equally, it optimises the entry of DNAH2, a core component of the axoneme motility apparatus. In the absence of CEP76, the sperm axoneme is functionally incompetent. Following development of the core axoneme, CEP76 is required for positioning of the annulus and thus influences mitochondrial sheath length. In addition to its TZ function, our data reveal that analogous to its roles in somatic cells [37], CEP76 plays a role in the suppression of centriole duplication in haploid male germ cells. This study highlights *CEP76* as a bona fide male fertility gene in men and mice and adds to the growing evidence that several genetic factors contribute to both sperm morphology and sperm count.

It has recently been suggested that the annulus and TZ are part of the same ciliary structure [29] and our data support this idea. The annulus shares a number of proteins with the TZ and the localisation of several of these proteins (e.g., CEP290, MKS1) follows annulus migration during late spermiogenesis [49]. Consistent with this, the abnormal annulus structures observed in sperm from *Cep76* knockout males suggests that TZ structure influences annulus structure and function. Specifically, we show that CEP76 is required for annulus positioning, and, in its absence, the annulus is poorly formed and migrates a shorter distance. Very little is known about factors that govern annulus migration and positioning [50], but these processes do not appear to be influenced by development of the fibrous sheath that ultimately borders the annulus in mature spermatids but is formed prior to annulus migration. This is highlighted by data from *Akap3* and *Akap4* knockout mice, which exhibit poor sperm fibrous sheath development, annulus positioning and midpiece length [20, 30, 51]. Equally, the absence of an annulus in *Sept4* and *12* mutant models did not impair fibrous sheath formation [28, 32].

It has been hypothesised that within CEP76 the C2 and transglutaminase domains cooperate to allow CEP76 to play a role in the transition zone as a Y-shaped linker [40]. Y-shape linker structures form the main body of the TZ; proteins of which can be grouped into the Meckel syndrome (MKS) and nephronophthisis (NPHP) complexes [52]. These complexes participate in multiple roles during ciliogenesis – TZ attachment, core axoneme extension and then regulation of components entry into the cilia/flagella [53]. Based on domain architecture and functional predictions, CEP76 has been hypothesised to span both MKS and NPHP complexes [40]. The presence of a C2 domain in CEP76 strongly suggests it anchors at the side of the Y-shaped linker interacting with the plasma membrane [40]. Our data highlight that, *in vivo*, CEP76 is not essential for axoneme formation, for example compared to CEP290, where CEP290 absence precludes TZ formation and ciliogenesis altogether [54, 55]. Rather, data suggest that CEP76 plays a facilitative role in TZ function specifically in male germ cells and is likely not a core Y-linker component. As mice did not show symptoms of MKS or NPHP, our data reveal CEP76 is not essential for somatic cells TZ function. The possibility exists, however, that mice possessed subtle defects in somatic tissues.

Further, our proteomics results revealed the content of only a small number of sperm tail were altered in the absence of CEP76, supporting the hypothesis that CEP76 is highly selective in its function and that other TZ proteins are required to form a functioning sperm tail. This is consistent with the concept that TZ content is cell type-specific and that different TZ proteins are involved in the selective recruitment of precise subsets of proteins from the many thousands of proteins present with the cell cytoplasm, i.e., the TZ functions as a cell type-specific filter controlling entry into cilia.

Overt consequences of CEP76 loss included the abnormal assembly and content of sperm tail accessory structures. We observed inappropriate packaging of the fibrous sheath and the absence of spacing between the circumferential ribs. AKAP4 is the most abundant fibrous sheath protein, constituting nearly half its entire content [56], and acts as a scaffold protein for both the longitudinal columns and ribs, while AKAP3 plays a similar role in the ribs of the fibrous sheath [21, 57]. The fibrous sheath defects seen in the absence of CEP76 are consistent with the poor entry of major fibrous sheath proteins into the sperm tail compartment via the TZ. Similarly, we identified an accumulation of granulated bodies in the sperm neck region and the partial absence of outer dense fibres in the principal piece axoneme. Collectively, our results strongly suggest the deficit in the entry of fibrous sheath and outer dense fibre components into the tail compartment limited the normal formation of accessory structures and thus directly impaired sperm motility.

Finally, we identified significant aggregation of mitochondria and morphology in the midpiece. The mitochondrial sheath is the last accessory structure to be loaded onto the sperm tail. This occurs in parallel with the migration of the annulus and plasma membrane down to a position immediately proximal, and abutting, the principal piece, as marked by the start of the fibrous sheath, [24, 58]. Although the membrane migration occurs at the same time as annulus migration, it does not appear to be driven by the annulus as the former occurs even when the annulus does not form [28, 30]. While the annulus is not required for mitochondrial sheath formation, in its absence mitochondrial morphology is frequently abnormal [28, 31, 59]. Prior to their loading, spherical mitochondria are recruited from the cytoplasm and ordered in four columns parallel to the axoneme. They then move towards the core of the tail and attach to the outer dense fibers, elongate, coil and stagger, to intercalate around the midpiece [58]. Next, mitochondria elongate and attach end-to-end to form a double helix around the axoneme. *Cep76* knockout sperm mitochondria were largely elongated and uniform along the midpiece, suggesting that mitochondrial recruitment and early mitochondrial elongation processes proceeded normally. We predict the shorter annulus migration distance in sperm from *Cep76* knockout males leaves insufficient space for the normal number of mitochondria to coil around the sperm tail, i.e. sterically interfere with the later processes of mitochondrial elongation and prevent their tight compaction within the midpiece [60]. Ultimately, the patency of the annulus to act as a barrier between major sperm tail compartments also likely influenced the loading of mitochondria, as demonstrated by examples of mitochondria in the principal piece section.

Our data revealed a second function of CEP76 in male germ cells – a role in supressing supernumerary centrioles, analogous to its role identified in human sarcoma cells *in vitro* [37]. A previously established mechanism highlighted that CEP76 controls centriole duplication via interaction with PLK1 and CP110 [36, 37]. While not investigated directly here, we note *Cep76* knockout mice were viable and were free of any overt body wide disease, suggesting that CEP76 is not essential for centriole function in somatic tissues.

To the best of our knowledge, this is the first example of a protein with a germ cell-specific function in the TZ. In the absence of CEP76, essential components of the sperm tail are unable to enter the ciliary lobe, meaning less or minimal incorporation into the growing tail and thus male infertility. These data provide support for the concept that TZ composition is cell type-specific and that the TZ provides an additional layer of specificity to the composition and function of cilia and flagella.

## Materials and methods

### Ethics statement

Experimental procedures involving mice followed animal ethics guidelines generated by the Australian National Health and Medical Research Council (NHMRC). All animal experiments were approved by the Animal Experimentation Ethics Committee (BSCI/2017/31) at Monash University, or The University of Melbourne Animal Ethics Committee (application 20640).

### Knockout mouse production

As described previously [34], exome sequencing of infertile men identified *CEP76* as a high confidence candidate male fertility gene. The patient carried a homozygous c.607G>C (p.Gly203Arg) missense variant in a conserved residue of *CEP76*, which was predicted to be intolerant to variation (Figure S1). Pathogenicity assessments using SIFT and Polyphen-2 [61, 62] predicted the missense variant to affect function (SIFT) and to be possibly damaging (Polyphen). To test the requirement for CEP76 in male fertility, *Cep76* knockout mice were generated on the C57BL/6J background through the Monash University Genome Modification Platform (a partner of the Australian Phenomics Network) using CRISPR/Cas9 technology. Excision of exon 3 of *Cep76* was undertaken with CRISPR guide sequences targeting regions flanking exon 3: upstream – TTTAAAACTCAGTTCGTGGT and downstream –GGTCTACATAGTAAAGTTCT. This was predicted to lead to a premature stop codon in exon 4 and a truncated protein in the only protein-coding transcript of *Cep76* (ENSMUST00000097542.3). Changes in gene sequence were identified with Sanger sequencing. Mice heterozygous for the *Cep76* deletion were intercrossed to generate knockout individuals and wild type controls. Genotyping was performed by Transnetyx (Corvoda, USA) using the primers F – CCCATTAACAGCCTCTGCTTCATAA and R – GAGACAGGGTTTCTCTGTGAATTCT. A reduction in *Cep76* transcript level (primers F – GCGGCTCGATTTGTTAATGT and R – AGTCCCCACACAGACAAAGG), was verified via qPCR on testis cDNA relative to *Ppia* (primers F – CAGTGCTCAGAGCTCGAAAGTTT and R – ACCCTGGACATGAATCCT).

### CEP76 species alignments

CEP76 protein alignments were conducted using the protein Basic Local Alignment Search Tool (NCBI). Sequences used were *Homo sapiens* ENSP00000262127; *Pan troglodytes* ENSPTRP00000016812; *Macaca mulatta* ENSMMUP00000069818; *Rattus norvegicus* ENSRNOP00000034590; *Mus musculus* ENSMUSP00000095149; *Danio rerio* ENSDARP00000075595. We compared entire protein identity across species and focused on the conservation of amino acid 203G, which was mutated in the infertile patient (Figure S1B).

### Analysis of *Cep76* expression

Whole organ RNA was extracted from adult mouse brain, epididymis, heart, liver, lung, spleen and testis, as well as testes from mice aged day 0-50, to investigate the expression of *Cep76* across different tissues and throughout the establishment of the first wave of spermatogenesis. Each tissue was homogenized in TRIzol Reagent to isolate RNA, which was converted to cDNA and used for qPCR with SYBR Green master mix as previously described [63]. Primers used to detect *Cep76* were F – CTCGGTCACCAGCAATGAAA and R – CAGACAGTGGTGAGGCCAAG, and housekeeping gene *Ppia* are denoted above. We also utilised single cell RNA sequencing data contained within FertilityOnline (https://mcg.ustc.edu.cn/bsc/spermgenes2.0/index.html) to investigate *Cep76* expression in mouse testes.

### Fertility analysis

Knockout males (*Cep76*^-/-^) and wild type male littermates (*Cep76^+/+^)* were aged to 10-14 weeks and their fertility was assessed using the pipeline outlined previously [64]. In brief, 5 mice of each genotype were setup to mate with 2 independent wild type females each (6-12 weeks old). The presence of a copulatory plug was recorded as an indication of successful mating. Litter sizes were recorded as the number of pups generated per plug. Males were subsequently culled and weighed (10-14 weeks of age), and one testis and epididymis were dissected and processed for histological assessment. Additional testes and epididymides were snap frozen on dry ice for calculation of daily sperm production and epididymal sperm counts as described previously [65]. In addition, sperm were collected from the cauda of the epididymis through backflushing, then resuspended in MT6 medium at 37°C for motility assessment via computer assisted semen analysis [66]. Residual sperm were washed in PBS then dried onto SuperFrost slides overnight. Sperm were then fixed in 4% paraformaldehyde for 10 min and washed in PBS and stained with Mayer’s haematoxylin for 10 min and eosin (Amber Scientific, Midvale, Australia) for 1 min to allow an assessment of sperm morphology and tail length. Alternatively, fixed sperm were permeabilized in 0.1% Triton-X-100/PBS (Sigma Aldrich, Castle Hill, Australia) for 10 min, washed in PBS and stained with 10 µg/ml DAPI (ThermoFisher Scientific, Scoresby, Australia) to allow an assessment of sperm head morphology, or 1 μg/ml peanut agglutinin conjugated to AlexaFluor-488 to label the acrosome. Head morphology assessment was performed using Nuclear Morphology Analysis software version 1.17.1 [67] via an ImageJ plugin (National Institutes of Health, USA). All length measurements (midpiece length, tail length and annulus positioning) were measured using ImageJ (version 1.52k).

### Electron microscopy

To investigate germ cell ultrastructure, testes were processed for electron microscopy as outlined previously [68]. Similarly, caudal sperm were backflushed into MT6 medium and processed for electron microscopy as outlined previously [68]. Images were taken either on a Jeol 1400 Plus electron microscope at the Vera and Clive Ramaciotti Centre for Electron Microscopy (Monash University, Australia), or a Talos L120C or a FEI Teneo VolumeScope at the Ian Holmes Imaging Center (The University of Melbourne). To view the mitochondria, annulus, and fibrous sheath structure of sperm via SEM, sperm were isolated from the cauda epididymis and incubated in 100 µl of 1 x PBS for 30 min to strip the plasma membrane then processed as outlined in [69].

### Scoring of mitochondrial sheath, fibrous sheath and annulus normality

To quantify the degree of ultrastructural defects within sperm tail structures, scoring of each structure (mitochondrial sheath, fibrous sheath, and annulus) was performed on SEM images. For each biological replicate, ten sperm were assessed. Mitochondrial sheath normality was scored 1-5: 1 – missing mitochondria; 2 – many abnormally oriented and/or thick mitochondria; 3 – few abnormally oriented and/or thick mitochondria; 4 – broadly normal with very few abnormally oriented mitochondria; 5 – no defects where ‘normal’ was defined as homogeneously coiled mitochondria, with none missing, thick or abnormally oriented. Fibrous sheath normality was score from ranks 1-3: 1 – no slits and/or massive aggregation of circumferential ribs or longitudinal columns; 2 – reduced number of slits or slight aggregation; 3 – normal structure, where normal was defined as slits occurring at regular intervals and no aggregation. Annulus normality was score as: normal – intact, not shrunken, and localised to the junction between the midpiece and principal piece; or abnormal – small, ectopically placed or unidentifiable.

### Mass spectrometry to define sperm protein composition

Sperm were backflushed from caudae epididymides of adult wild type (1 mouse per replicate, *n* = 3 replicates) and *Cep76* knockout males (2 mice per replicate due to reduction in epididymal sperm content) into MT6 medium for 15 min at 37°C. Sperm were washed three times in 1 x Tris buffered saline, pelleted, and then stored at -80°C. Sperm pellets were then dried in a speed vacuum and prepared for liquid chromatography tandem mass spectrometry and run on a SCIEX QTRAP6500 as described previously [70].

Data were assessed as 1) total/raw spectral counts or 2) total spectral counts were normalised to the alpha tubulin content (alpha tubulin chain 8) for a comparison of sperm tail protein content in recognition of the shorter sperm tails measured from *Cep76* null mice. Sperm tail proteins were identified via proteomics or localisation studies. For completeness, proteins with an unknown localisation or that are localised throughout the head and tail were included in the tail protein group. Proteins associated exclusively with the sperm head or neck and mitochondrial proteins were not included here as the normalisation process (to alpha tubulin) as they are not core sperm tail proteins. A two-tail t-test was used to determine significant differences in protein content between genotypes. All data are available in Table S1.

### Immunofluorescence and protein localisation

To define centriole number in male germ cells, spermatids were isolated from the testes of males using the STAPUT method [71]. Cells were settled onto poly-L-lysine coated SuperFrost slides for 15 min and then fixed in ice cold methanol at -20 °C for a maximum of 7 min. Slides were immediately washed and rehydrated in PBS. Fixed spermatids were permeabilised in 0.2% Triton-X-100/PBS for 10 min, blocked in CAS-block (Dako), incubated overnight in 0.1 µg/ml Centrin antibody (Merck 04-1624) to stain centriole components and beta tubulin (Abcam ab21057) to stain the manchette for spermatid staging, then counterstained with DAPI. Z-stacks were taken using a Leica SP8 confocal and flattened to capture all slices in a single frame. The number of centriole components per spermatid was then counted for 100 cells per replicate, per genotype.

To define the localisation of a sub-set of differentially expressed proteins, fixed sperm were permeabilised in 0.2% Triton-X-100/PBS, blocked in CAS-block, incubated overnight in primary antibodies (0.5 µg/ml DNAH2 [Invitrogen 64309], 2.5 µg/ml SEPT4 [Abcam 166788]) at 4 °C, stained with relevant fluorescent secondary antibodies (ThermoFisher Scientific) for 1 h, then counterstained with DAPI. For other antibodies (2.8 µg/ml AKAP4 [ProteinTech 24986-1-AP], 0.8 µg/ml SUN5 [ProteinTech 17495-1-AP]) sperm were permeabilised in 0.5% Triton-X-100/PBS. For the assessment of sperm mitochondrial sheath length, live sperm were loaded with 5 µM Mitotracker Red CMXRos in MT6 solution for 30 min as per the manufacturer’s instructions. Sperm were then washed in PBS, fixed in 4% paraformaldehyde and allowed to settle onto slides, as described above. All images were taken using cellSens software (Olympus, Notting Hill, Australia) on an Olympus BX-53 microscope (Olympus) equipped with an Olympus 392 DP80 camera or a Leica SP8 confocal. Immunofluorescence staining intensity was used as a surrogate of protein content in sperm tails. FIJI v2.1.0 was used to trace sperm tails (freehand line tool) and measure the average pixel intensity for AKAP4 and DNAH2 staining, to account for differences in tail length.

CEP76 localisation in testis sections, germ cells and sperm was attempted with two antibodies: Abcam ab86613 and Bethyl #A302-326A. Both antibodies reacted with antigens in knockout tissue.

### Statistical analysis

Statistical analyses to determine significance between wild type and *Cep76* knockout data were performed in Prism 9 (GraphPad, San Diego, USA) using a student’s t-test on average data per mouse. A nested t-test was used to analyse sperm midpiece length, sperm tail length, annulus migration distance, mitochondrial sheath and fibrous sheath ultrastructure score, and pixel intensity analysis of protein entry in the sperm tail. In these datasets, multiple sperm were measured or assessed within a single biological replicate and each data point was used. A p value less than 0.05 was considered significant.

## Author roles

BH, AEOC, JM, GS, JD and MB undertook experiments.

BH, GS and MKOB analysed data.

BH, JD and MKOB designed the experiments and wrote the paper.

All authors provided intellectual input and feedback on drafts.

All authors reviewed and approved the final publication.

**Table 1 A. Proteins with significantly altered content in sperm from *Cep76* knockouts.**Green proteins are upregulated in sperm from knockout mice, blue proteins are downregulated.

**Table 1B. Proteins with significantly altered content in sperm from *Cep76* knockouts, after normalisation to alpha tubulin content.** All proteins were upregulated in sperm from knockout. Known sperm tail proteins are highlighted orange. Other proteins are present throughout the entire sperm cell or localisation is unknown.

---

**Supplementary Figure 1.**
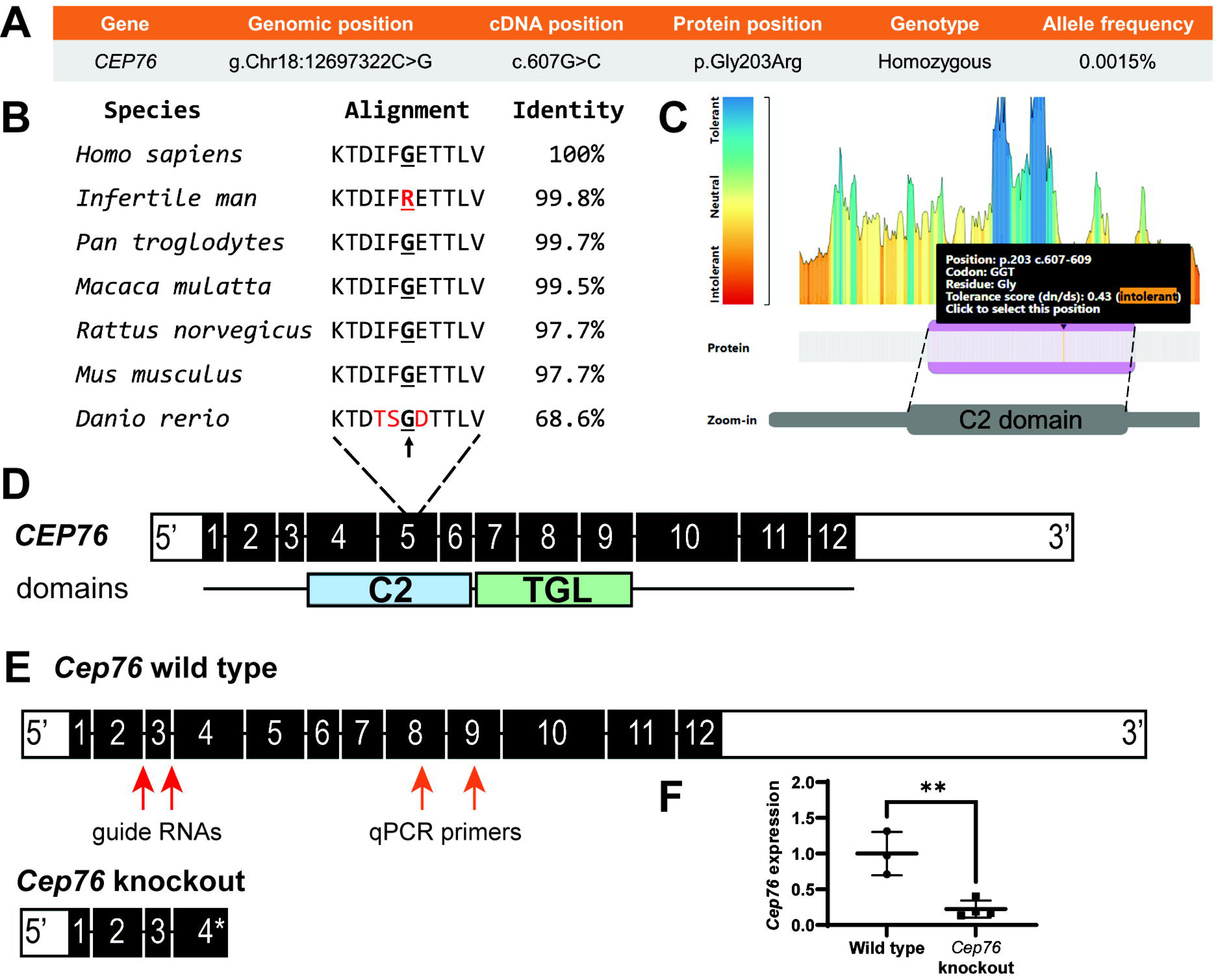
*CEP76* human genetic variant and *Cep76* knockout mouse model. A. CEP76 mutation identified in an infertile man, including genetic coordinates and protein position. B. CEP76 protein species alignment, including conserved amino acid (G, bolded + underlined), and total protein identity across species. Red letters denote amino acids not conserved in zebrafish (*Danio rerio*) and the affected amino acid in the infertile man. C. MetaDome assessment of the affected amino acid and its tolerant to change, which was assessed as intolerant. D. The genetic variant affected exon 5 of the human CEP76 protein, within the C2 (ciliary targeting, exons 4-6) domain. CEP76 additionally holds a TGL (transglutaminase, exons 7-9) domain. E. Mouse full length *Cep76* transcript and *Cep76* knockout transcript shown below (* denotes premature stop codon in exon 4). Red arrows denote where guide RNAs for exon 3 removal targeted – intronic regions surrounding exon 3; orange arrows denote approximate target sites of qPCR primers. F. *Cep76* expression as measured by qPCR in wild type and knockout testes, relative to housekeeper *Ppia* expression. ** = *p* < 0.01.

**Supplementary Figure 2.**
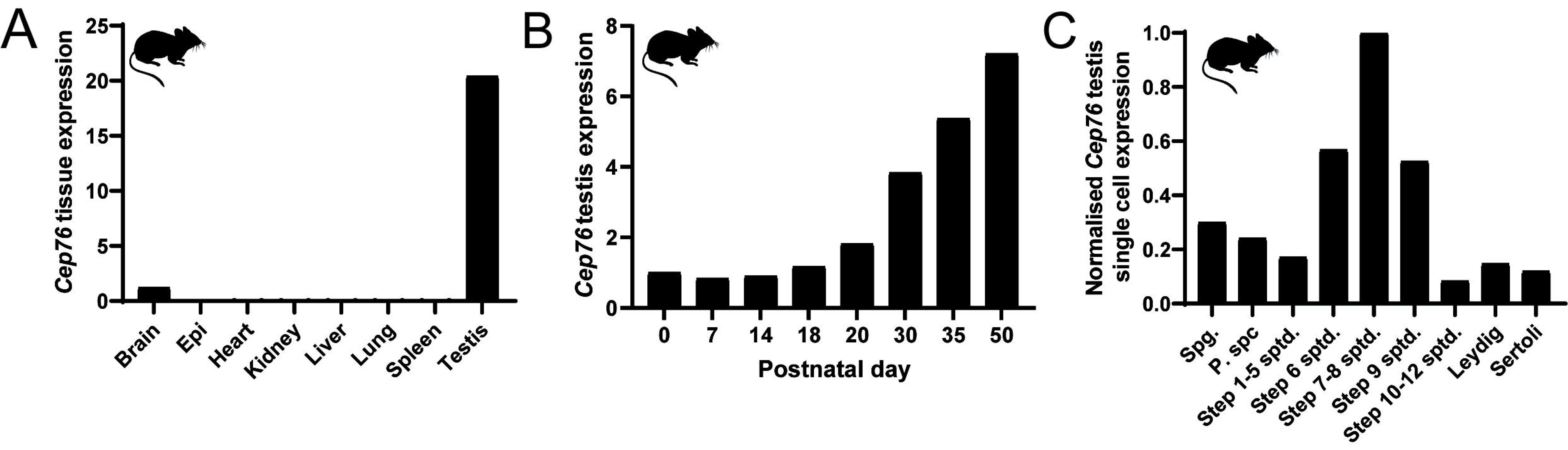
*Cep76* is a spermatid enriched gene. A. *Cep76* expression across major organs in mice as assessed by quantitative PCR (qPCR) relative to housekeeper *Ppia*. B. *Cep76* expression in mouse testes during the first wave of spermatogenesis, relative to *Ppia*. Ages correspond to the first appearance of different male germ cell types. C. *Cep76* expression in mouse testis cell types as defined by single cell sequencing. Spg. = spermatogonia, P spc. = pachytene spermatocyte, sptd. = spermatid.

**Supplementary Figure 3.**
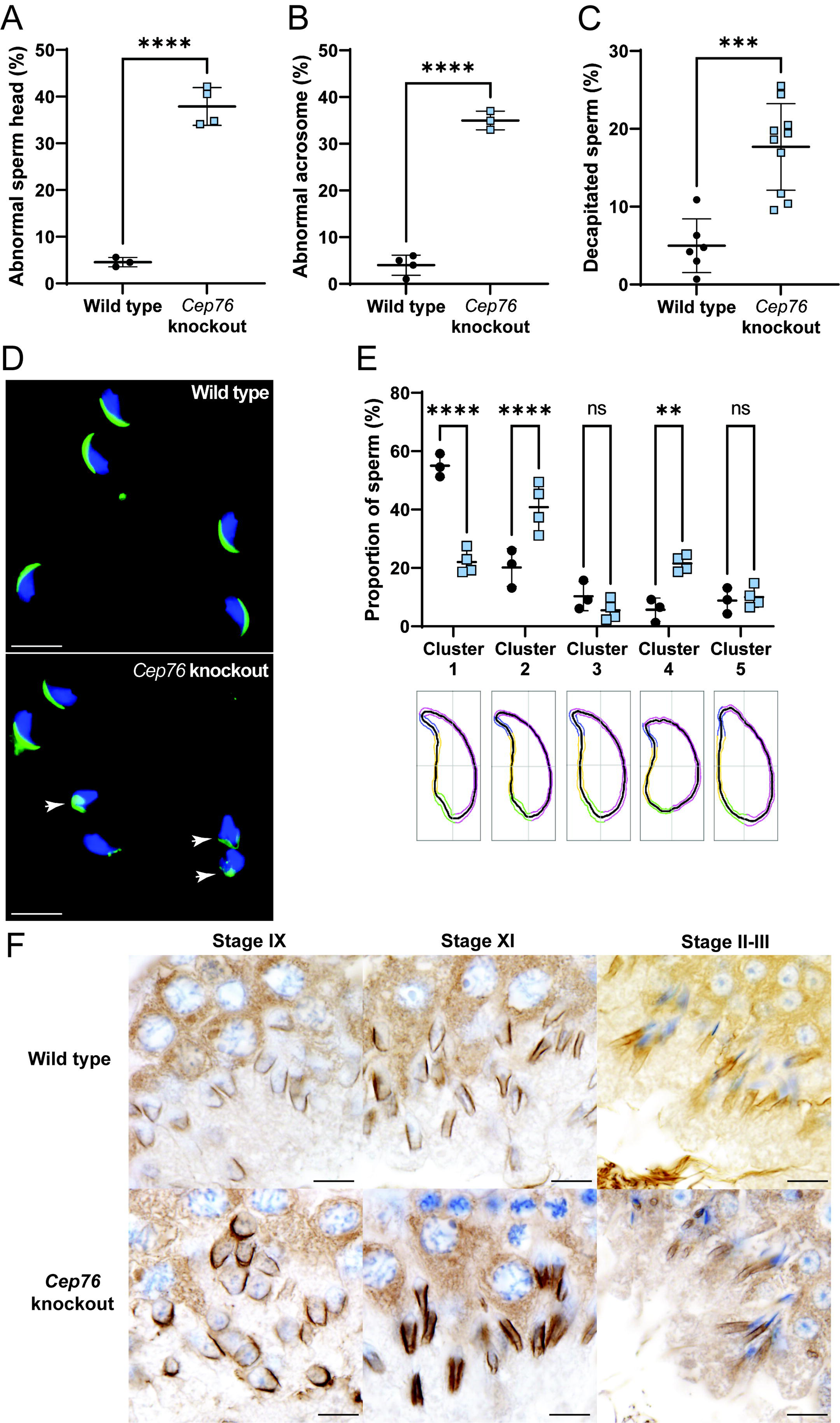
CEP76 is required for sperm head and acrosome shape but not normal manchette formation. A. Abnormal sperm head shape assessment. B. Abnormal sperm acrosome formation assessment. C. Sperm decapitation assessment. D. Sperm head and acrosome morphology assessment on cells stained with DAPI and PNA. E. Objective sperm head shape analysis and the proportion of sperm falling into each Cluster. Cluster 1 are most normal through to Cluster 5 as the most abnormal head shapes. Representative traces are shown for head shapes below each cluster. F. Spermatids at stage IX, XI and II-III stained with alpha-tubulin to mark the manchette are shown. Scale bars are 10 µm in length. ** *p* < 0.01, **** *p* < 0.0001.

**Supplementary Figure 4.**
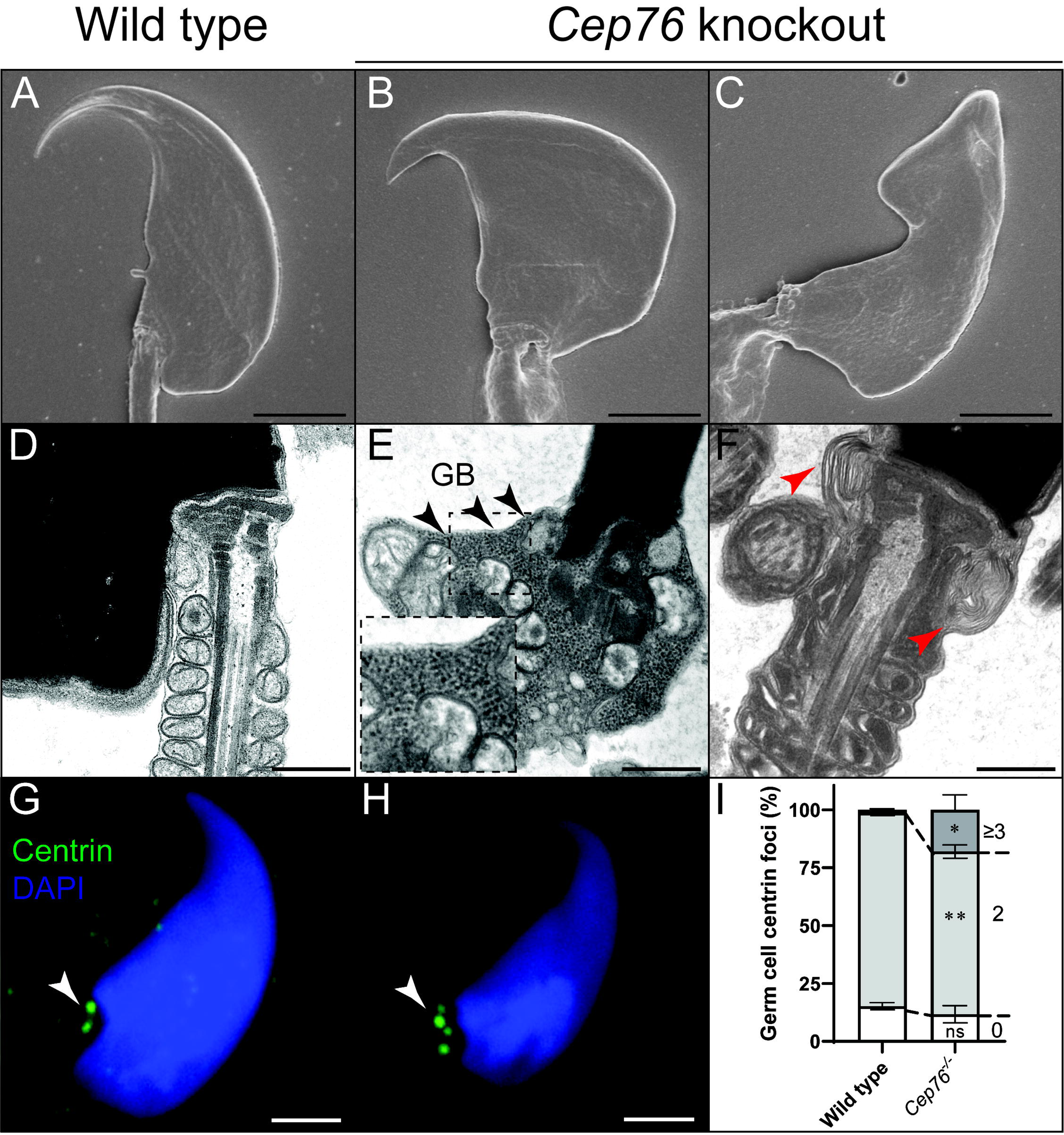
CEP76 is required for sperm neck integrity and suppression of centriole duplication in male germ cells. A-C. Scanning electron microscopy images of sperm heads from wild type and *Cep76* knockout sperm. D-F. Transmission electron microscopy images of sperm neck regions. Arrows and expanded dashed box denote potential granulated bodies (GB) throughout the neck region in panel E. In panel F, the red arrows point to abnormal membrane folding. G, H. Centriole number/content as defined by centrin staining. Arrows point to individual centriole components. Scale bars = 2 µm in all panels, except length in D-F where they are 1 µm. I. Quantification of centriole number in round and elongating spermatids, categories as 0, 2 or 3+. * *p* < 0.05, ** *p* < 0.01, ns = not significant.

